# Disturbance of the Sense of Agency in Obsessive-Compulsive Disorder and its Modulation by Social Context

**DOI:** 10.1101/2024.06.27.600968

**Authors:** Manuel J. Roth, Axel Lindner, Andreas J. Fallgatter, Andreas Wittorf, Aiste Jusyte

## Abstract

Executing precise actions and perceiving them as one’s own is a fundamental ability underlying the sense of agency (SoA). The SoA thereby heavily relies on the accuracy and reliability of forward models, capturing sensory movement consequences. Impairments thereof thus represent a promising candidate mechanism contributing to cases of SoA pathogenesis. In obsessive-compulsive disorder (OCD), for example, the feeling of control over one’s actions is perturbed: Compulsive actions are often experienced as uncontrollable and performed without conscious awareness. At the same time, compulsions can be coupled with an inflated sense of illusory control for uncontrollable events. Here we studied self-action perception in virtual reality with and without veridical or rotated visual feedback about subjects’ pointing movements to test whether patients’ internal forward models are indeed less reliable compared to controls. Interestingly, OCD patients did not exhibit deficits in their accuracy and reliability of motor performance and self-action perception in the absence of visual feedback, suggesting intact forward models. Nonetheless, OCD patients weighted rotated visual action-feedback significantly stronger perceptually. Furthermore, they adapted their movement to this false feedback on a trial-by-trial basis. Finally, increasing the social relevance of action consequences led to stronger feedback weighting in all participants while this effect increased with the strength of OCD symptomatology under conditions with strongest social relevance. We suggest that internal forward models are equally reliable in OCD but their weight is pathologically decreased leading to patients’ overreliance on explicit visual action-feedback and, more generally, to their over-attribution of unrelated events to themselves.

## Introduction

The ability to experience authorship, i.e., the feeling of control over one’s own actions and, through those actions, also over one’s environment, is not only a prerequisite for a coherent sense of self and a sense of responsibility and self-efficacy. It is also of major importance for learning and the pursuit of individual goals - and thus for development. This experience has been coined Sense of Agency (SoA, (Haggard 2017)). Given the fundamental role of SoA in perception and action, studying altered SoA and its association with psychopathology can be a promising avenue for elucidating the mechanisms underlying complex psychiatric conditions. One such condition is obsessive-compulsive disorder (OCD). OCD is a difficult to treat condition with a lifetime prevalence of 1%-2% (Fawcett, Power et al. 2020) that emerges early in development and is associated with high levels of chronicity (Wewetzer, Jans et al. 2001). Symptoms of OCD can be divided into two broad clusters: obsessions, i.e. intrusive unwanted thoughts, and compulsions, i.e. stereotypical repeated actions/routines (e.g., (Association 2013)). Previous research has conclusively linked OCD to an inflated sense of responsibility for uncontrollable negative events and moral sensitivity (Shapiro and Stewart 2011, Harrison, Pujol et al. 2012, Vicario 2012, Mancini and Gangemi 2015). Furthermore, patients often experience intense fear to cause harm to themselves or others and employ compulsory behaviors to “neutralize” these fears (Lopatka and Rachman 1995, Arntz, Voncken et al. 2007, Doron, Sar-El et al. 2012, D’Olimpio and Mancini 2014), which indicates an inflated illusory belief of control. Compulsive actions, however, are often experienced as compelling and uncontrollable, are performed without conscious awareness, and, after years of repetition, often become decoupled from the initial obsessive thoughts or a sense of relief (Reed 1983, Van Schalkwyk, Bhalla et al. 2016, Ferreira, Yucel et al. 2017). Also, 60-70% of OCD patients report that the repetitive behaviors are in fact driven by aversive sensory sensations that an action has been executed “incompletely” or “not just right”, rather than by specific fears (Ferrão, Shavitt et al. 2012). The execution of compulsions is thus not always tied to specific fears or aimed at neutralizing them, but appears to be a result from perturbed sensorimotor processing (Szalai 2019).

Taken together, the core symptoms of OCD appear to reflect a disturbed SoA. Moreover, as we will point out in the following, this disturbance could relate to different levels of the SoA. Crucially, the construction of a SoA comprises both predictive (sensorimotor) and reconstructive components (Haggard 2005, Moretto, Walsh et al. 2011, Synofzik, Vosgerau et al. 2013). The former builds on forward models of motor commands that allow to predict the sensory consequences of actions. These predictions can be used to discriminate between self-caused and non-self-caused sensory information (Blakemore, Wolpert et al. 2000, Frith, Blakemore et al. 2000, Lindner, Thier et al. 2005, Synofzik, Lindner et al. 2008, Synofzik, Vosgerau et al. 2009). Investigations into the role of such sensorimotor predictions based on forward models as low-level correlates of agency have thus been frequently employed, emphasizing their importance in constructing a reliable SoA (Lindner, Thier et al. 2005, Haggard and Tsakiris 2009, Synofzik, Thier et al. 2010, Gentsch and Schütz-Bosbach 2011, Kühn, Nenchev et al. 2011, Gentsch, Kathmann et al. 2012, Waszak, Cardoso-Leite et al. 2012, Wilke, Synofzik et al. 2013, Weller, Schwarz et al. 2017). The reconstructive components of SoA, on the other hand, reflect the fact that other information, such as retrospective inferences triggered by sensory action outcomes, is also integrated into the experience of agency (Moore and Obhi 2012, Gentsch and Synofzik 2014). In fact, existing evidence shows that the valence of movement outcomes influences the experience of agency, for example, in temporal binding (Moretto, Walsh et al. 2011, Takahata, Takahashi et al. 2012, Yoshie and Haggard 2013, Beck, Di Costa et al. 2017, Borhani, Beck et al. 2017, Di Costa, Théro et al. 2017, Yoshie and Haggard 2017, Tanaka and Kawabata 2019), sensory attenuation (Hughes and Waszak 2014, Hughes 2015), and the self-attribution of sensory action-consequences (Wilke, Synofzik et al. 2012).

In the case of OCD, the impaired SoA may, at first glance, be more likely to concern this second, reconstructive level, since patients exhibit an overall inflated sense of responsibility and control. However, whether the weight of this reconstructive process is pathologically increased in OCD or, alternatively, whether this increased weight reflects a compensation of defects on the predictive/sensorimotor level is still open. Existing evidence so far cannot reliably answer whether the deficit lies at the predictive or reconstructive level or at both.

A study by Gentsch and colleagues that examined the difference of a neurophysiological signal, the N1 component in EEG, in response to a self-generated sensory stimulus versus a non-self-generated stimulus, found that this difference was significantly smaller in OCD patients. The authors therefore proposed a dysfunctional forward model mechanism in OCD that results in imprecise sensory predictions to account for this difference between patients and controls (Gentsch, Schütz-Bosbach et al. 2012). According to current accounts of perceived agency, such reduced reliability of internal (e.g., sensorimotor signals, predictions, intentions) agency cues would lead to a stronger relative weighting of other (e.g., explicit feedback, valence of action outcomes) cues in the construction of a SoA (Synofzik, Vosgerau et al. 2009, Gentsch and Synofzik 2014, Fradkin, Adams et al. 2020). Existing evidence indeed shows that subclinical analog groups with high OC tendencies (Lazarov, Dar et al. 2010, Lazarov, Dar et al. 2012, Ezrati, Friedman et al. 2019) and patients with OCD (Lazarov, Liberman et al. 2014) are more susceptible to the effects of false feedback. In line with these findings, and based on the fact that doubt and uncertainty are a central feature of OCD, Lazarov and colleagues proposed the Seeking Proxies for Internal States (SPIS) model (Lazarov, Liberman et al. 2014, Lazarov, Liberman et al. 2014, Dar, Lazarov et al. 2021). This model suggests that OCD can be characterized by attenuated access to internal states, i.e., states that cannot be reliably assessed by outside observers, such as physiological states, intentions, etc., for which patients try to compensate by relying (more) on other proxies, such as rules, rituals, explicit feedback, i.e., any information that seems less ambiguous. The authors and colleagues present studies that seem to support this notion, showing, for example, impaired performance in producing certain muscle tensions and a greater tendency to demand/rely on explicit proxies when available, albeit most of the evidence comes from non-clinical populations (Lazarov, Dar et al. 2010, Lazarov, Dar et al. 2012, Lazarov, Dar et al. 2012, Lazarov, Liberman et al. 2014, Lazarov, Cohen et al. 2015, Dar, Eden et al. 2019, Ezrati, Friedman et al. 2019).

Here we aimed to test both the predictive and the reconstructive components of agency in patients with OCD compared to healthy controls. To this end, we employed a manual pointing paradigm in virtual reality that allowed us to provide veridical (0°) or rotated (±10°, ±30°) visual feedback on participants’ pointing movements, or no visual feedback at all. This well-established method allows to compare the actual, the fed-back, and the subjectively estimated movement direction, and thus to measure the weight each participant gives to the (rotated) visual feedback relative to their actual movement. Also, we estimated participants precision in pointing towards an explicit visual target without feedback. With respect to OCD, however, one could argue that previous studies on agency lack similarity to real-world scenarios in which compulsions are often carried out specifically with the intent to prevent negative outcomes for oneself or others. Therefore, we wanted to characterized the contributions of external agency cues like social context that tap into the inflated sense of responsibility in OCD. Specifically, we investigated how outcome valence (monetary gains and losses associated to one’s actions) and outcome reference (i.e., whether consequences of an action affect oneself or others) influences participants actions and the perception thereof.

To address this, we included experimental manipulations of outcome valence and social relevance by implementing conditions that included monetary gains and losses, either for the participant or for a charitable organization. Using this design, hence we can determine (I) the precision/reliability of participants’ movements, (II) the precision/reliability of their sensory predictions concerning these movements, and thus the precision of self-action perception, (III) the impact of (possibly false) visual feedback on (I) and (II), and, finally, the influence of social and valence cues on these measures.

We expect to find a stronger reliance of patients’ self-action estimates on visual feedback, when provided. If this stronger reliance were based on imprecise signals describing the sensory effects of own movements, as suggested in the study by Gentsch (Gentsch, Schütz-Bosbach et al. 2012), this should be apparent in more noisy estimates regarding the sensory consequences of own movements in patients compared to controls. Our hypothesis concerning the manipulation of outcome valence and social context (gaining/losing money for oneself or a charitable organization) was that patients with OCD are more strongly influenced by negative outcome valence and especially if others are affected by it. This should be reflected in our measures of self-action and self-action perception. Given our hypotheses of impaired SoA in OCD in general, we furthermore expected to find a correlation between OCD symptom strength and the amount of visual feedback weighting and especially in cases when the outcome of an action has a consequence for others.

## Methods and Materials

### Sample and Procedure

Inpatients and Outpatients currently receiving treatment for OCD were recruited from the University Hospital for Psychiatry and Psychotherapy, Tübingen, Germany. Inclusion criteria for the patient sample were: 18 – 65 years of age, no history of schizophrenia spectrum disorders, no current use of sedating medications or substance abuse, normal or corrected-to-normal vision. A group of healthy controls with no current psychopathology was recruited according to the same inclusion criteria as the patient group. The recruitment was carried out via the university mailing list. Ultimately, 39 patients with OCD and 40 healthy controls participated in our study. We excluded one patient (substance abuse) and one healthy control (eating disorder), leaving a group of 38 patients (12 males, mean age 35 years) and 39 controls (13 males, mean age 32 years) for the final analysis (Supplementary Table S1). All participants were right-handed. The project was approved by the Ethics Committee of the University Hospital of Tübingen. All participants gave written informed consent and received monetary compensation.

### Clinical and control measures

Interested participants were screened for exclusion criteria by telephone and, if eligible, were invited to participate in various assessments, including self-report measures, a clinical interview conducted by a trained clinician, and the actual experiment. The DSM-5 diagnostic criteria for OCD as well as diagnoses that warrant exclusion were assessed using the Mini International Neuropsychiatric Interview for DSM-5 (American Psychiatric Association 2013). OCD symptom severity was assessed using the Yale-Brown Obsessive-Compulsive Scale (Goodman, Price et al. 1989) and the Obsessive-Compulsive Inventory Revised (OCI-R) (Foa, Kozak et al. 1998). The Obsessive-Compulsive Trait Core Dimension Questionnaire (OCTCDQ) was used to assess feelings of incompleteness and harm avoidance related to obsessions or compulsions (Summerfeldt, Kloosterman et al. 2001).

**Table 1.** Sample Demographics. Sex and age. distributions were not significantly different between groups (sex: t(75) = 0.1622, p = .872; age: t(75) = −1.5926, p = 0.115). Patients scored significantly higher on all dimensional measures of OCD symptoms (OCI-R: t(75) = −12.6121, p < 0.001; OCTCDQ: t(75) = −11.6761, p < 0.001). Detailed participant information is provided in Supplementary Table S1. CTR: Control group; OCD: Obsessive-Compulsive Disorder; OCI-R: Obsessive-Compulsive Inventory-Revised, OCTCDQ: Obsessive-Compulsive Trait Core Dimensions Questionnaire; Y-BOCS: Yale-Brown Obsessive-Compulsive Scale

| Group | N | Males/Females | Age (M ± SD) | OCI-R (M ± SD) | OCTCDQ (M ± SD) | Y-BOCS (M ± SD) |
| --- | --- | --- | --- | --- | --- | --- |
| CTR | 39 | 13/26 | 32 ± 11 | 4 ± 2 | 5 ± 4 | NA |
| OCD | 38 | 12/26 | 36 ± 12 | 25 ± 10 | 40 ± 18 | 22 ± 8 |

### Experimental tasks

Participants performed pointing movements with their right index finger in a 2D virtual reality setup in which they could not see their own hand, but could receive visual feedback about their movement on a monitor viewed through a mirror (Figure 1a). Hand movements were recorded using a touchpad (Magic Touch USB; Keytec, Inc.). At the beginning of each trial, participants placed their index finger in the central starting position using a tactile cue on the surface of the touchpad. This resulted in the appearance of a white central starting point and a surrounding white target circle. After the target circle disappeared, participants performed a simple out-and-back pointing movement from the central starting point to a position in the upper right quadrant of the previously displayed target circle (radius 9 cm). Importantly, for the trial to be successful, the turning point of the movement had to be within a turning range around the target circle (± 1.5 cm), otherwise the trial was aborted and the participants received the feedback that the movement was too short/long. Similarly, movements that started before the disappearance of the target circle resulted in a trial abort. Apart from these constraints, the movement had to consist of one continuous motion and participants were asked to move quickly and smoothly. All aborted trials were repeated until the planned number of trials was successfully completed. After each trial, regardless of the type of trial (see below), participants estimated their pointing direction by placing a cursor in that direction (*Estimated Pointing Direction; EPD*). More precisely, after the movement was completed, a white dot appeared at a random position halfway between the starting point and the target circle and participants could move that dot along an angular axis using a trackball. They confirmed their decision with a mouse click. The overall experiment consisted of three experimental blocks described below, the order of which was pseudorandomized and counterbalanced between groups (patients, controls) to control for sequence effects.

**Figure 1.**
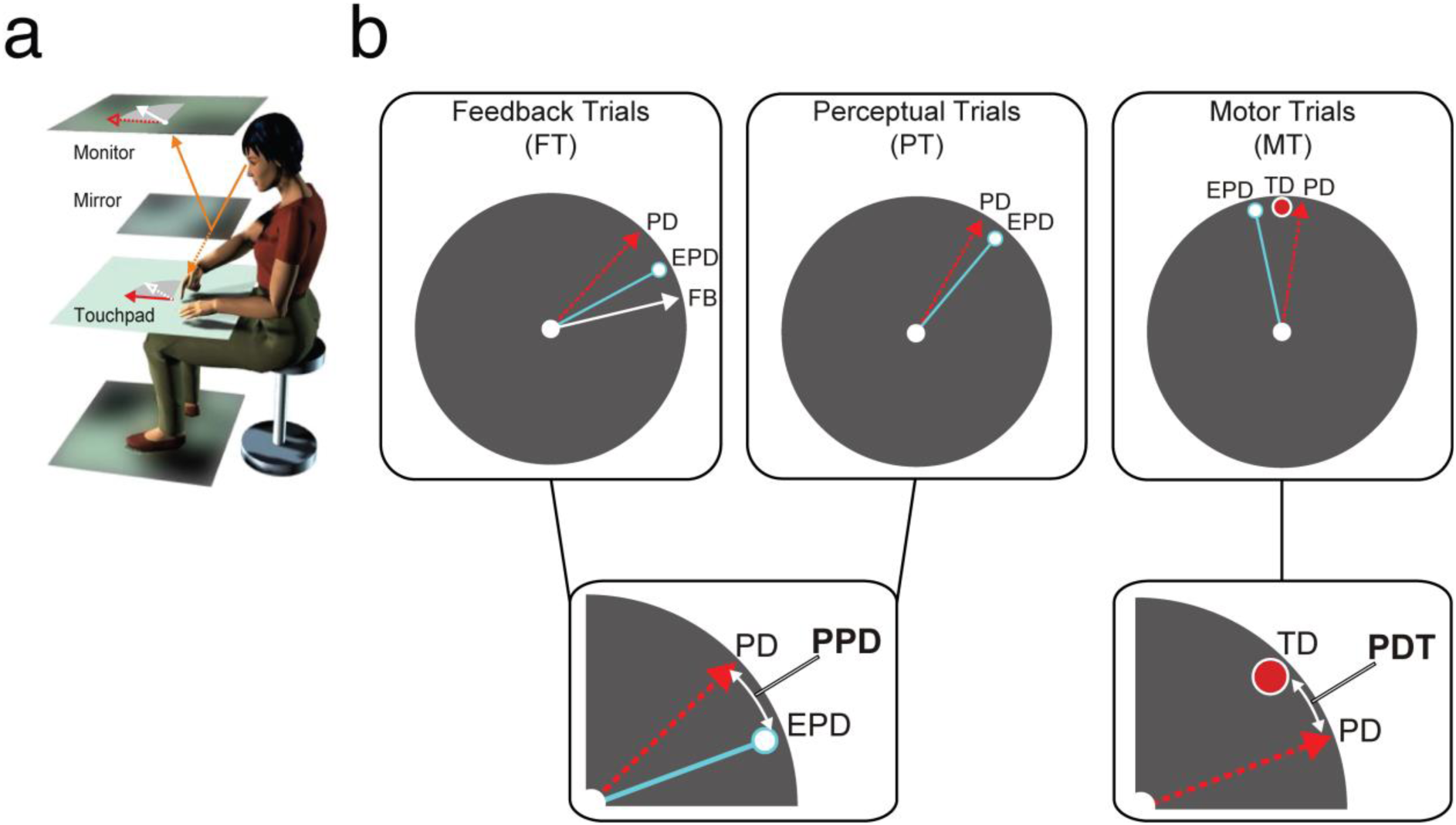
Experimental Setup and Design. a) Participants performed out-and-back pointing movements (red) on the table plane within the upper right quadrant (i.e., between 12 and 3 o’clock) of a presented target circle without seeing their hand. In each trial, they viewed a virtual image of the table plane and, depending on the current trial, their finger (white) on the feedback monitor via a mirror (solid orange). For geometric reasons, the virtual image appeared in the same plane as the subjects’ finger movements (dotted orange). b) At the end of each trial, participants reported their estimated pointing direction (EPD, light blue) by placing a cursor in the direction of the movement performed. In feedback trials (FT), participants performed pointing movements in a freely chosen direction and received feedback about their movement. The feedback (FB, white) could be either veridical or rotated by ±10° or ±30° relative to the true pointing direction (PD, red). In perceptual trials (PT), participants performed pointing movements in a freely chosen direction (PD, red) without any feedback. In motor trials (MT), a visual target was presented at one of four possible positions (TP, red disk with white border) to which participants had to point without any feedback. PD: Pointing Direction, EPD: Estimated Pointing Direction, FB: Feedback, PPD: Perceived Pointing Direction = EPD-PD, TD: Target Direction, PDT: Pointing Direction relative to the Target = PD- TD.

#### Baseline Block

The *baseline block* consisted of three different randomly interleaved types of trials: *motor trials (MT)*, *perceptual trials (PT)*, and *feedback trails (FT)*.

During the MT, participants saw a visual target (red dot) appear randomly at one of four possible positions (equal share), namely at 12, 1, 2, or 3 o’clock relative to a clock face, in the upper right quadrant of the target circle before the start of the movement. This visual target then disappeared along with the target circle, providing the go-signal. Participants were asked to point to this target without any visual feedback about their movement.

During PT, participants could point in any direction within the upper right quadrant of the target circle, but were asked to distribute their movements evenly across this area throughout the whole baseline block. They did not receive any visual feedback about their movement.

In FT, participants were again free to choose their pointing direction, but received visual feedback about their movement (a white disc representing the participant’s fingertip) throughout. Importantly, the visual feedback could be either veridical (α = 0°) or be rotated by ±10° or ±30° around the starting point of the movement and relative to the participant’s true pointing direction. The feedback rotation in the current trial was randomly selected and the five possible rotations were equally distributed across all FT.

A successful trial (i.e., straight motion, turning point within the turning range, etc., see above) was indicated by a change of the feedback cursor changing to a larger yellow circle in the FT, or the starting point changing to a larger yellow circle in the MT and PT. Additionally, a beep sound (1000 or 3000 Hz, randomly selected) was presented. As described above, after successfully completing the pointing movement, participants were asked to estimate their true pointing direction (EPD) in all types of trials. The baseline block consisted of a total of 134 trials and included 24 MT, 30 PT, and 80 FT presented randomly interleaved.

#### Self and Other Blocks

The *self* and *other blocks* included only PT and FT. Both blocks included a valence manipulation with monetary gains and losses on each trial. Participants were informed that their performance would determine the outcome, i.e., either a reward or a deduction of money (average amount per trial: 4 ± 0.8 Euro Cents), without disclosing any determinant of success. In reality, gains and losses were randomly distributed across trials and kept constant between participants (such that each participant won approximately 2.50 Euros per block). During a trial, current wins/losses were indicated by a change of the feedback cursor (FT) or the starting point (PT) into a larger green (gain) or red (loss) circle with the won/lost amount displayed inside. In addition, one of the two different sounds indicated a gain (3000 Hz) or a loss (1000 Hz). Wins and losses were summed across all trials, and the current total was displayed after each trial. This valence manipulation served to investigate whether the experience of agency for rewards and punishments is modulated by self-reference of these effects in OCD. To manipulate self-reference, all participants were informed that the additional monetary gains would be paid either to the participant (self block) or to a charitable organization (other block). In the case of the other block, participants chose one of four real-world charities (World Wildlife Fund, Medecins Sans Frontieres, UN Refugee Agency, UNICEF) at the beginning of the block. At the end of the experiment, the participants were informed that they had been deceived and that all the money would actually be paid out to the participants. Apart from these manipulations, the general task and trial structure was the same as in the baseline block. The self- and other blocks consisted of a total of 110 trials, each comprising 30 PT and 80 FT presented randomly interleaved.

### Analyses

#### Movement Data

All analyses were performed using MATLAB R2018b (The MathWorks, Inc.). The data was stored for offline analysis. We defined the direction of pointing movements by a straight line fitted to the position samples of the outward part of the movement using linear regression analysis. MT with a deviation in movement direction of more than 40° relative to the overall median were discarded. The same was true for trials with an excessively curved trajectory (movement deviating from a straight line by more than 20% of the instructed movement amplitude). To eliminate possible sampling artifacts, we also removed trials with peak velocities above 100°/s. The number of excluded trials was small, and an average of 97% (FT), 98% (MT), and 97% (PT) of the trials was used for analysis.

#### Measures of Motor Performance and Self-Action Perception

To characterize and compare the accuracy and reliability of participants’ movements and their perception of these movements, we focused on two main variables, PDT and PPD (see Figure 1b). The variable describing participants’ motor accuracy, as measured during MT in the baseline block, was their true pointing direction relative to the target. Therefore, we compared participants’ true pointing direction (PD) with the target direction (TD) of the visual target by subtracting TD from PD. We refer to this variable as *PDT*, i.e., *Pointing Direction relative to Target*: PDT = PD - TD. The smaller the absolute value of this variable, the better participants were at hitting the target. On the other hand, in order to have a measure of participants’ self-action perception, we compared participants pointing direction (PD) with their estimated pointing direction (EPD) by subtracting PD from EPD. We refer to this variable as *PPD, i.e., Perceived Pointing Direction*: PPD = EPD - PD. The PPD provides us a measure of how accurately participants perceive their own movement. More importantly, the PPD allows us to estimate how much the rotated visual feedback influences participants’ perception, as indicated by a shift in PPD toward the direction of the rotated feedback and away from their true PD (i.e., larger PPD compared to veridical feedback). For both PDT and PPD, we estimated their median value across trials for each individual participant. In addition to these measures of accuracy, we also characterized the precision of participants’ actions and action-perceptions by estimating their respective reliabilities. We define reliability as the inverse of each individual’s variance (1/s^2^) in the respective measure (PDT, PPD). When analyzing the FT, we applied an offset correction by subtracting the value at the 0° rotation (i.e., trials with veridical feedback) from all other rotations to minimize interindividual variance (Wilke, Synofzik et al. 2013). For more information see Supplemental Results and Figure S1. When comparing the influence of feedback rotation on the PPD across groups, we focus in particular on the larger, suprathreshold rotations of ±30° (Wilke, Synofzik et al. 2013). To study the influence of such large rotations independent of the direction of rotation, we constructed a mean for PPD and PDT between - 30° and 30° while using the inverse of the negative rotation (otherwise the effects of the positive and negative rotations would cancel each other out). Finally, to analyze the difference between outcomes, i.e., gain/loss of money during the self and other blocks, across all three blocks, trials in the baseline block were selected based on the tone presented (high or low). Since there was no difference in outcome associated with tone type in the baseline block, the effect of each tone should not be different and therefore serves as a good control measure.

#### Statistical Analysis

Statistical analyses were performed using Matlab R2018b (The MathWorks, Inc.) and SPSS 27 (IBM). Participants’ performance was analyzed and compared by means of ANOVAs. We used Mauchly’s test to test for sphericity and adjusted the F statistic using Huynh-Feldt-correction when the assumption of sphericity was not met. For the correlation analyses, we tested for normally distributed residuals in linear correlation analysis. Since the tested residuals were normally distributed, we resorted to Pearson correlation. To control for multiple comparisons, we applied Bonferroni correction when necessary (see main text).

## Results

### Baseline Block

We first compared the performance of both groups’ in MT to ensure that any group differences in later analyses were not due to the patients’ inability to produce accurate pointing movements in the context of our experimental setup. In MT, participants had to point to a presented visual target without any visual feedback. The mean deviations of participants’ movements from the pointing targets (PDT) were small and not statistically different between groups (patients: *M* = 0.96°, *SD* = 3.25; controls: *M* = −0.08°, *SD* = 4.35, *t*(75) = −1.177, *p* = .243), indicating accurate performance in both groups (Figure 2a). The same was true for movement reliability (see Methods), which was not statistically different between the two groups (Figure 2b, patients: *M* = .05, *SD* = .029; controls: *M* = 0.056, *SD* = 0.043, *t*(75) = 0.685, *p* = 0.495).

**Figure 2.**
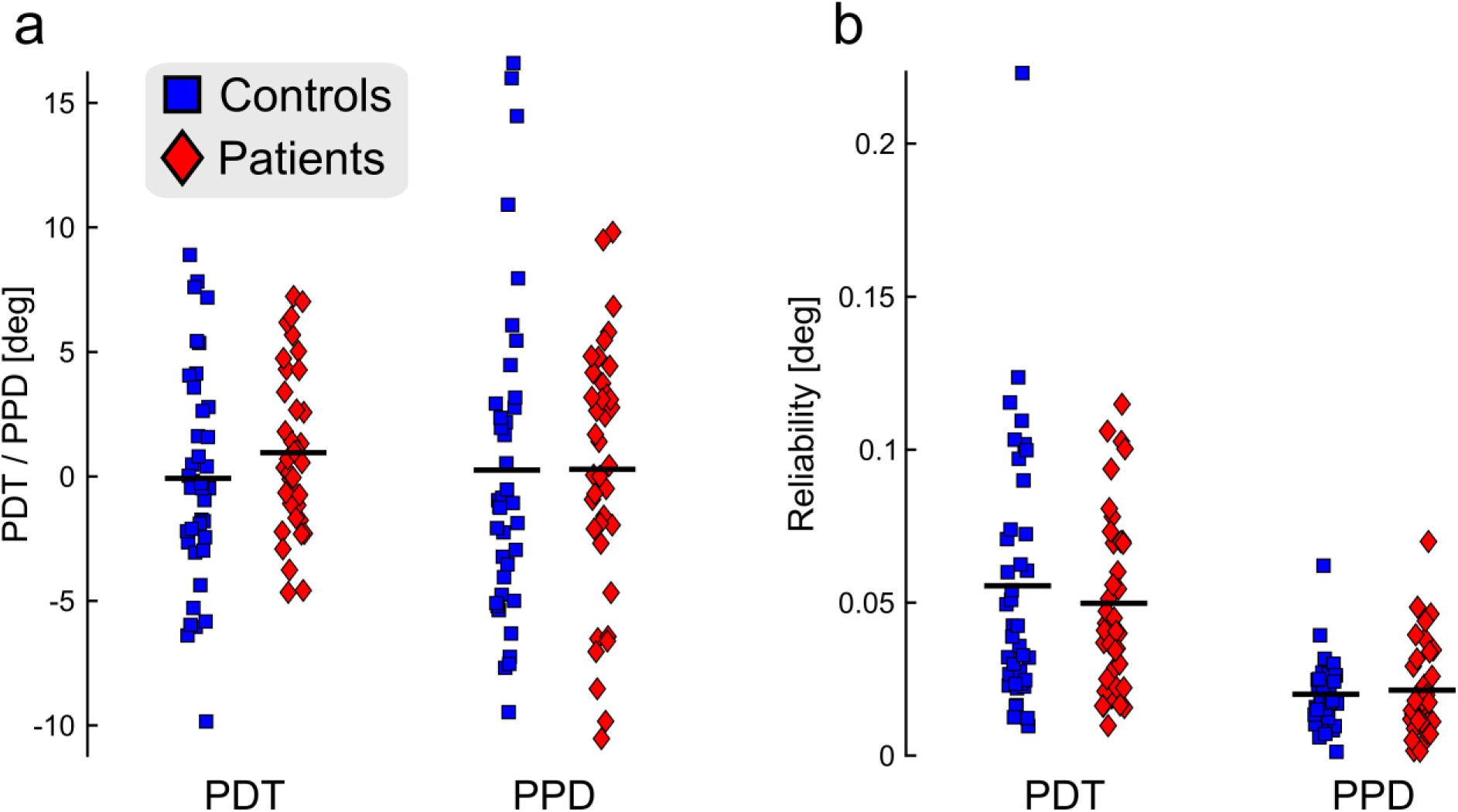
Accurate and reliable performance in both groups during motor trials (MT) and perceptual trials (PT). a) In MT, participants had to point to a visual target without feedback about their movement. The mean pointing direction relative to the target (PDT) of both groups was close to zero and did not differ between groups. In PT, participants performed pointing movements in a freely chosen direction without any visual feedback. Mean perceived pointing directions (PPD) were similar and not statistically different between groups. b) The reliability (1/s^2^) of the participants’ PDT and PPD was likewise similar in both groups. Thus, both groups were equally able to execute movements with comparable accuracy and reliability. Individual data points represent the median of each participant, thick black lines represent the group mean.

Next, we analyzed whether self-action perception per se was impaired in patients in the baseline block. We compared perceived pointing direction (PPD) in PT, i.e., trials in which participants made pointing movements in a freely chosen direction without visual feedback. PPD was not significantly different between patients and controls (*t*(75) = -0.024, *p* = 0.981; Figure 2a). Again, both groups were highly accurate, with mean deviations of their estimated pointing directions from the true pointing direction amounting to 0.26 ± 6.4° for controls and 0.29 ± 5.1° for patients. The reliability of the participants’ PPD (Figure 2b) was likewise not different between the two groups (*t*(75) = -0.454, *p* = 0.651; patients: 0.021 ± 0.015, controls: 0.02 ± 0.011).

In summary, the motor performance of patients in our task did not differ from that of healthy controls, neither in the movements themselves nor in the perception of these movements. Thus, possible agency distortions in OCD were not attributable to imprecise or inaccurate motor output (PDT) or self-action perception (PPD) per se, the latter often being interpreted as a proxy for the accuracy of forward models.

Importantly, according to our hypotheses we expected to find evidence for increased weighting of false visual feedback in the patient compared to the control group, as a reflection of the agency distortions described in the introduction. Therefore, we analyzed participants’ perceptual judgments in FT, trials with veridical or rotated visual feedback, which allow to estimate the extent to which participants’ PPD is influenced by visual feedback distortions. Results from FT are shown in Figure 3a. Feedback rotation had a clear influence on participants’ perception, which was confirmed by a significant main effect of rotation (*F*(1.54, 115.5) = 9.057, *p* < 0.001) in a 4 x 2 mixed-model ANOVA with the factors Rotation (±10, 30°) and Group. As expected, we also found a significant Rotation x Group interaction (*F*(3, 75) = 3.679, *p* = 0.013), which was likely driven be the obvious group difference at suprathreshold rotation of ±30°. In order to analyze this effect more closely, we compared the average PPD at 30° rotation directions (using the inverse of the negative rotation, see Methods) between groups with a one-way ANOVA. This revealed significantly stronger weighting of visual feedback in patients (*F*(1, 75) = 4.739, *p* = 0.033); also see baseline block in Figure 4). In conclusion, OCD patients’ perception of their own movements was significantly more susceptible to strong visual feedback distortions compared to healthy controls. Thus, they appear to weight this feedback information more heavily relative to other internal movement-related information, such as sensory predictions, etc. Interestingly, this apparent lower relative weighting of internal agency cues was not due to lower precision or accuracy of these cues, which did not differ from controls (compare above and Figure 2). Across the overall sample of subjects, however, did the reliability of the PPD actually determine the weight of the visual feedback, as suggested in the Introduction? We indeed found that PPD reliability was significantly negatively correlated with the strength of feedback weighting at 30° rotation, such that the noisier participants’ PPDs were, the stronger the influence of the visual feedback on these estimates (compare Supplemental Figure S2).

**Figure 3.**
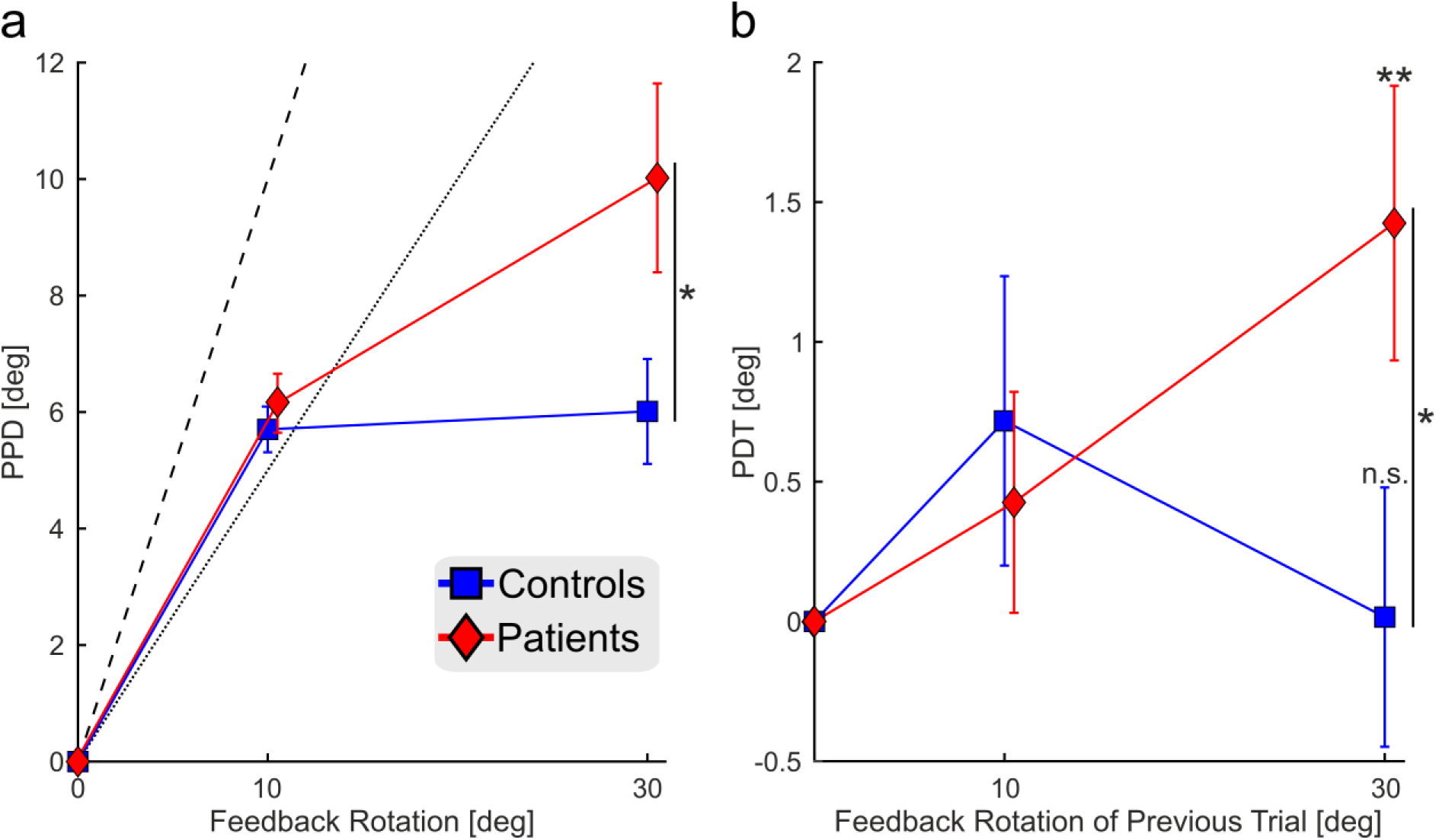
Influence of rotated visual feedback on participants perception and pointing accuracy (mean ± sem). a) Perceived pointing direction (PPD) in feedback trials (FT). The ordinate represents the magnitude of the PPD in degrees. The actual rotation of the visual feedback is plotted on the abscissa. An average PPD of zero represents the true pointing direction, i.e., participants’ perceptual estimates reflect only their actual movement, while the dotted and dashed lines reflect the perceptual weight of the rotated visual feedback, i.e., how much participants’ perceptual estimates are influenced by the false visual feedback by 50% (dotted) or 100% (dashed), respectively. There was a clear influence of feedback rotation on participants’ PPD, as indicated by a shift of PPDs away from zero. While the relative weight of visual feedback was 50% or more in trials with 10° rotation, it decreased in trials with 30° rotation, a finding consistent with previous studies (e.g., (Wilke, Synofzik et al. 2013)). In the case of 10° rotation, the results of both groups largely overlap, whereas they differ in the case of 30° rotation trials. More precisely, the patients’ perception is significantly more biased towards the 30° rotated visual feedback compared to the controls. b) Pointing direction relative to the visual target (PDT) in motor trials (MT). Patients with OCD showed a clear adaptive response in their pointing accuracy to a visual target in MT when the trial was immediately preceded by a FT with 30° rotated visual feedback. This adaptation was significantly stronger than in the control group, in which this effect was completely absent. Thus, the influence of exaggerated weighting of false visual feedback in patients extends to subsequent movements, even though the feedback is not systematically rotated and no gain in performance or reward is associated with adapting to it. * = p < 0.05; ** = p < 0.01; n.s. = not significant

**Figure 4.**
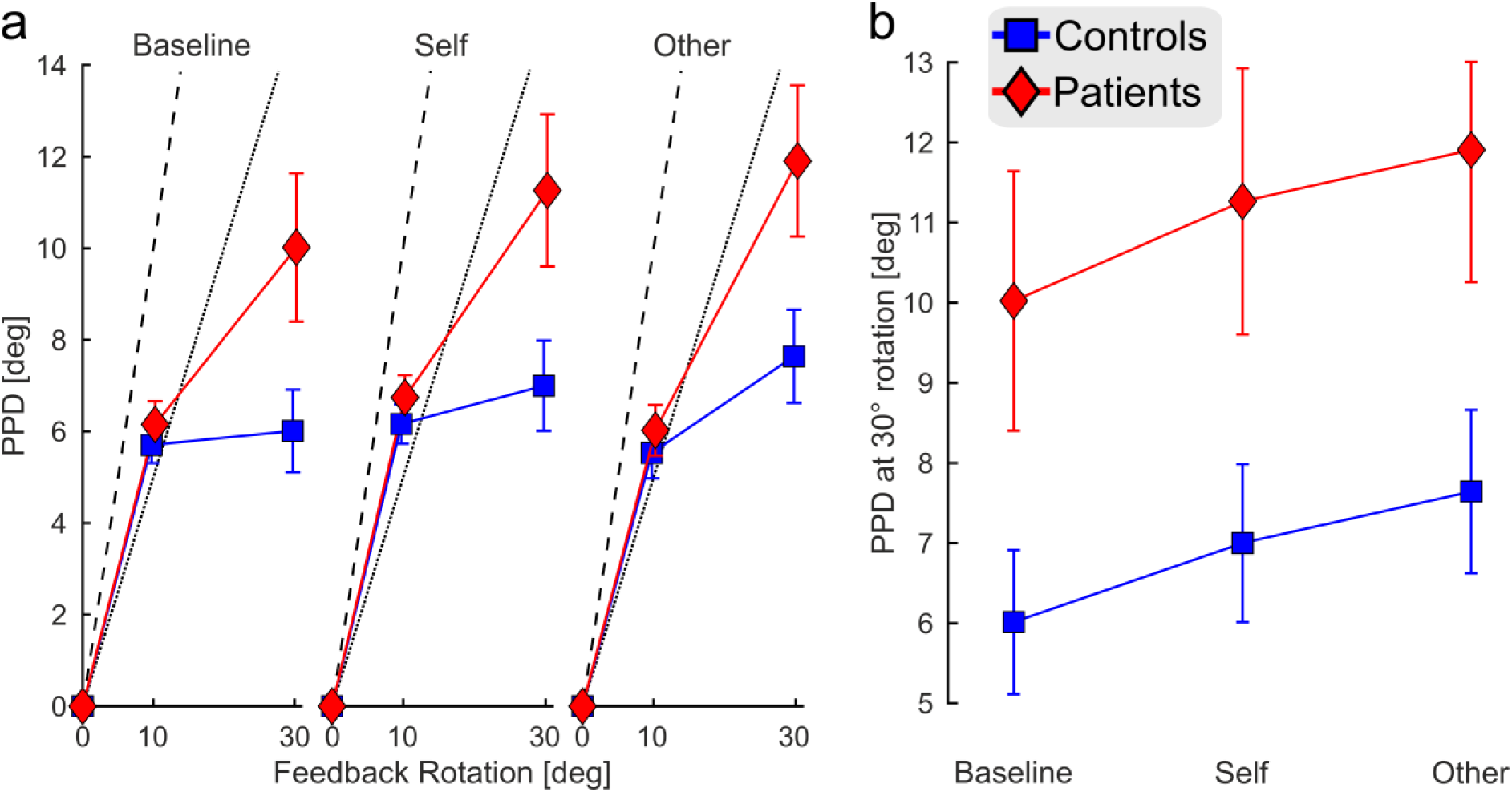
Results of feedback trials (FT) across experimental blocks and groups (mean ± sem). a) Perceived pointing direction (PPD) in feedback trials (FT) in the three different blocks. The ordinate represents the magnitude of the PPD in degrees. The actual rotation of the visual feedback is plotted on the abscissa. An average PPD of zero represents the true pointing direction, i.e., participants’ perceptual estimates reflect only their actual movement, while the dotted and dashed lines reflect the perceptual weight of the rotated visual feedback, i.e., how much participants’ perceptual estimates are influenced by the false visual feedback by 50% (dotted) or 100% (dashed), respectively. In all three blocks, feedback rotation had a clear influence on participants’ PPD, as indicated by a shift of PPDs away from zero, with largely overlapping results between patients and controls in trials with 10° rotation, but clear differences in the case of 30° rotation trials, with a significantly stronger bias towards the 30° rotated visual feedback in patients. b) Average PPDs in trials with 30° rotation across the three blocks for controls (blue) and patients with OCD (red). Patients show a significantly stronger weighting of visual feedback in their PPDs. This effect persists across all three experimental blocks. In addition, there was a significant increase in feedback weighting from the baseline to the self to the other block across participant groups.

Given this overreliance on rotated visual feedback during self-action perception in patients with OCD, we sought to examine whether this overreliance also biased patients’ movements in subsequent trials. In other words, given the strong influence of ±30° rotated visual feedback on patients’ PPD, one might expect to see traces of this effect on participants’ movements on a trial-by-trial basis. Specifically, right-rotated feedback should lead to pointing movements that are rotated to the left to compensate for (part of) this apparent error between intended and actual movement. We therefore analyzed the accuracy with which participants pointed to a visual target without visual feedback during MT. Importantly, we only included trials that were immediately preceded by FT, i.e., trials with visual feedback, and compared the mean (see Methods) PDT, i.e., our measure of pointing accuracy (Figure 3b). In line with the exaggerated weighting of ±30° rotated visual feedback in estimating pointing direction, patients not only subsequently adapted their movements in response to such trials (one-sample t-test; *t*(36) = 2.862, *p* = 0.007), but did so to a significantly greater extent than controls (independent-samples t-test, two-tailed, *t*(74) = −2.074, *p* = 0.042). Controls, on the other hand, showed no significant adaptation effects compared to trials following FT with veridical visual feedback (*t*(38) = 0.034, *p* = 0.973). Thus, the higher weighting of strongly rotated visual feedback in patients with OCD was not only evident in their estimated pointing direction, but it also had a direct influence on subsequent movements, despite the absence of any motor error or trial characteristics that would reward such an effect.

### Self- and Other Block

With the self- and other block, we investigated how outcome valence (monetary gains and losses) and outcome reference (i.e., whether the consequences of an action affect oneself or others) influence sensorimotor agency. We first analyzed the influence of block (baseline, self, other) and outcome (gains, losses) on PPD in PT (i.e., trials without visual feedback). The respective ANOVA revealed no significant main or interaction effects for the accuracy or reliability measure. Thus, there was no general difference between blocks or outcome valence in participants’ ability to perceive their own hand movements without visual feedback. As a next step we investigated PPD in FT (Figure 4a). According to our hypothesis, we expected to find differences in the influence of valence outcome and reference between patients and controls, namely that patients with OCD are more strongly influenced by negative outcome valence, especially if others are affected by it. Thus, we ran a 3 x 2 x 2 x 2 mixed-model ANOVA with the within-subjects factors block (baseline, self, other), outcome (gain, loss), and rotation (10°, 30°) and group (patients vs. controls) as a between-subjects factor on the average values of PPD. We found significant main effects of block (*F*(2, 150) = 6.319, *p* = 0.002) and rotation (*F*(1, 75) = 16.477, *p* < 0.001), and a group difference (*F*(1, 75) = 4.096, *p* = 0.047). In addition, there were significant interaction effects of rotation x group (*F*(1, 75) = 7.712, p = 0.007), block x rotation (*F*(1.07, 143.04) = 4.833, *p* = 0.01), and block x rotation x outcome (*F*(1.786, 133.956) = 4.425, *p* = 0.017). Therefore, contrary to our hypothesis, neither outcome valence nor reference had any effects related to group. However, there was an overall group difference, which is further supported by the rotation x group interaction since the difference between groups is, as predicted, most prominent at 30° rotations.

Thus, we next ran a post-hoc 3 x 2 x 2 mixed-model ANOVA with factors block (baseline, self, other), outcome (gain, loss), and group with the data of 30° rotation trials only, yielding the expected main effect of group (*F*(1, 75) = 5.702, *p* = 0.019) and block (*F*(2, 150) = 7.9756, *p* < 0.001), and an interaction effect of block x outcome (*F*(1.613, 121) = 5.243, *p* = 0.024). Post-hoc analyses concerning the interaction effect with outcome revealed that the outcome effect was based on an outcome difference in the baseline block, in which the two different sounds had no meaning in terms of valence. Therefore, we did not analyze this effect any further. The main effect of block, however, signifies an increase in feedback weighting from the baseline to the self to the other block across subject groups (Figure 4b). This is further supported by a significant linear effect of the factor block in the ANOVA (*F*(1, 75) = 13.865, *p* < 0.001). Three post-hoc 2 (block) x 2 (group) mixed-model ANOVAs comparing blocks revealed a significant main effect of block between the baseline and the self block and between the baseline and the other block (baseline vs. self: *F*(1, 75) = 4.064, *p* = 0.047; baseline vs. other: *F*(1, 75) = 8.577, *p* = 0.005; self vs. other: *F*(1, 75) = 1.154, *p* = 0.286).

In sum, the results pattern indicates that OCD patients showed significantly stronger weighting of rotated visual feedback compared to controls. Furthermore, our experimental blocks influenced the degree of feedback weighting in both subject groups.

### Feedback Weighting and Symptom Strength

We expected to find a relationship between OCD symptom strength and the extent of feedback weighting, predominantly in the other block. In order to analyze this effect, we performed an analysis on contrasting high and low OCD subgroups of patients according to their summed score in the OCI-R. According to our hypothesis, we expected to find that the patient subgroup with more severe symptoms (high OCD group) should be significantly more affected by the other block compared to the subgroup with less severe OCD symptomatology (low OCD group). A 3 x 2 mixed-model ANOVA with factors block and Subgroup revealed the expected significant block x group interaction (*F*(2, 36) = 4.249, *p* = 0.018). In order to make the two blocks that included valence conditions more comparable between the two groups, we looked at participants’ performance during FT in the self- and other block relative to baseline (baseline not different between subgroups, independent-samples t-test: *t*(36) = 0.796, *p* = 0.431). Figure 5 shows that the high OCD group exhibited significantly stronger weighting (Bonferroni corrected alpha) of 30° rotated visual Feedback in the other block relative to baseline (one-sample t-test; *t*(18) = 3.267, *p* = 0.004), while this effect was absent in the low OCD group (one-sample t-test; *t*(18)= 0.236, *p* = 0.816). Furthermore, high OCD patients’ perception of self-action was, as hypothesized, significantly more affected by the other block (independent-samples t-test, one-tailed, *t*(36) = 1.935, *p* = 0.0304) leading to stronger weighting of false visual feedback in this block compared to the other two. Dividing the patient group into two subgroups based on the summed score in the second questionnaire used, the OCTCDQ, led to the same qualitative results (see Supplementary Results and Figure S3). Thus, the more patients were currently affected by OCD symptoms the less they relied on internal movement information in the other block but rather relied on the strongly rotated visual feedback.

**Figure 5.**
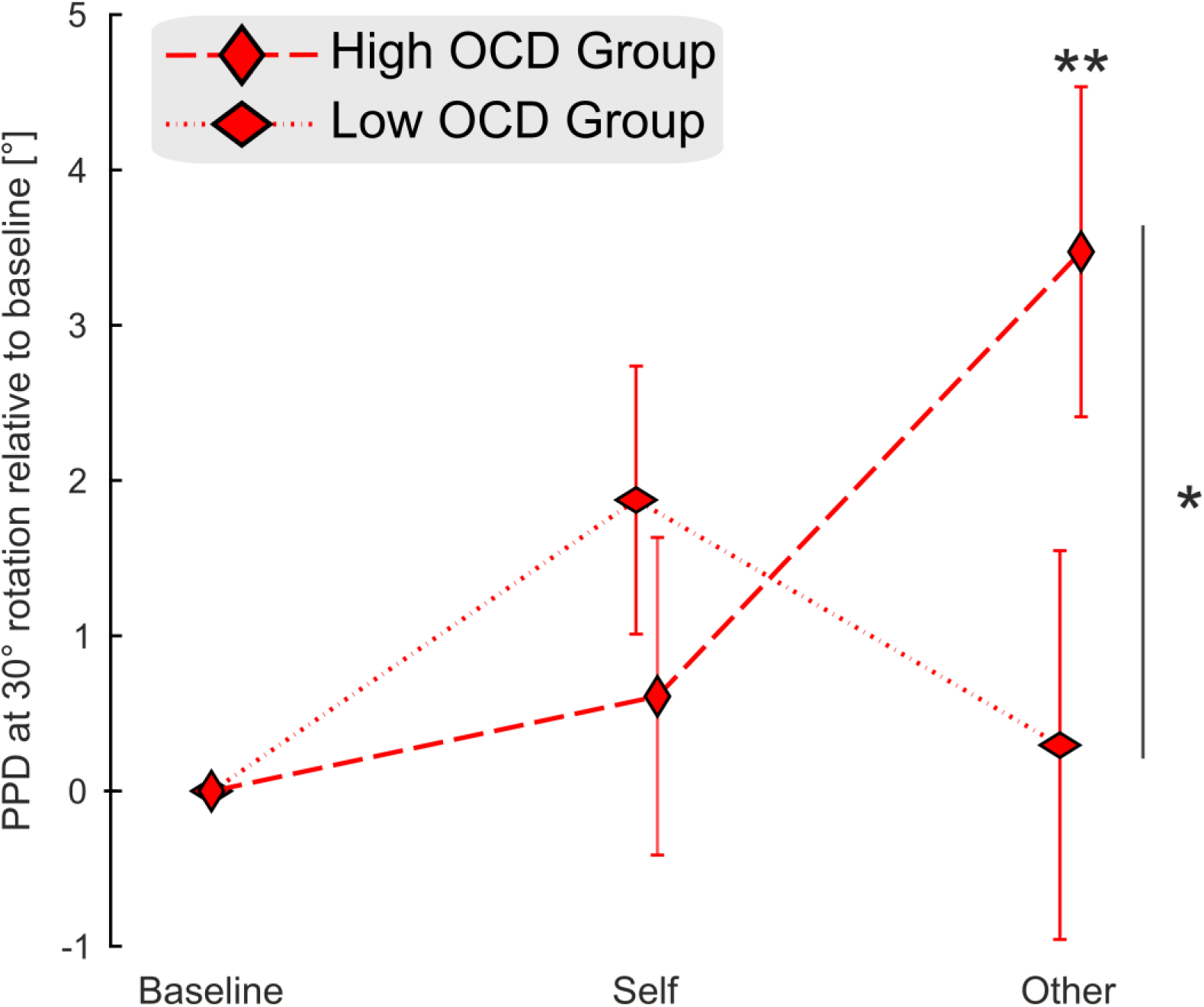
Differential effect of the experimental block on the two subgroups of patients depending on their symptom severity. Separating patients into two groups depending on their summed score in the OCI-R leads to clear differences in the other block. We expected this block to affect the high OCD group more strongly because it taps into the inflated sense of responsibility and moral sensitivity associated with the OCD symptomatology. * = p < 0.05; ** = p < 0.005.

## Discussion

The first aim of this study was to characterize sensorimotor components of agency in OCD. In line with our hypotheses, the results of our experiments showed that, compared to healthy controls, OCD patients exhibit excessive weighting of false visual feedback regarding their own hand movements. Thus, the integration of different agency cues, such as internal (i.e. resulting from execution of the movement itself) and external (e.g. the visual feedback) cue, was shifted towards false visual feedback. Furthermore, this shift in the integration of internal and external agency cues towards rotated visual feedback was not only visible in subjects’ self-action estimates but also directly affected patients’ subsequent movements. This was evident from a significant trial-by-trial adaptation to manipulated feedback, despite the absence of any overt motor error and the need to exhibit accurate movement. Importantly, none of these effects could be explained by imprecisions in movement or self-action perception *per se*, which did not differ between patients and controls. Thus, while patients’ motor as well as perceptual performance were both accurate and precise, they tended to over-rely on false visual feedback. A second aim of the study was to test whether social and valence aspects related to action outcomes can exacerbate the aberrant agency experience in patients with OCD. We found that social aspects, tested in two experimental blocks in which participants could gain/lose additional monetary rewards for themselves or a charitable foundation, had considerable influence on all participants’ perceptual estimations in FT, irrespective of subject group. More specifically, the influence of false visual feedback on participants’ estimates regarding their own hand movements was most pronounced under conditions when the social relevance was strong, i.e., when the action outcomes affected others. Furthermore, we found that OCD patients with stronger symptoms were significantly more affected by this condition compared to their counterparts with weaker symptomatology.

How could the overreliance on false visual feedback, both in perception and action, in OCD patients contribute to the described agency disturbances in this patient group? Previous research has highlighted the importance of attributing error information to their respective source in order to decide on how to react/adapt to it. Wilke and colleagues (Wilke, Synofzik et al. 2013) showed that during pointing, the magnitude of the visual error between actual and fed back movement direction determined whether this error was attributed to internal causes, i.e. whether it was self-caused, and consequently, how strongly participants adapted to it (also see (Wei and Kording 2009)). Conversely, the stronger weighting of false visual feedback and the trial-by-trial adaptation to said feedback in patients with OCD suggests that patients attributed a larger share of this erroneous feedback to themselves. Such an effect could explain many of the aforementioned irregularities in the SoA reported in OCD: Attributing “externally caused” erroneous information to oneself could be linked to an *inflated illusory belief of control* as well as to an *inflated sense of responsibility* (for actually uncontrollable (negative) events). At the same time, the attribution of unrelated errors to oneself would always lead to mismatches between accurately predicted and actually caused sensory afference. Hence, this may be a possible explanation why actions could often feel as “not just right” or “incomplete”, fitting the reports provided by patients with OCD. Further studies examining the link between the sensorimotor disturbances reported in this study and the clinical phenomena (not-just-right experiences, illusory control) are necessary in order to provide support for these possible explanations.

In general, constructing a SoA is a process in which one makes probabilistic inferences about current configurations of the environment and oneself acting on it. Therefore, different sources of information such as own actions, predicted sensory outcomes, and actual afferent sensory information, have to be integrated in a meaningful way. Following the influential predictive coding framework incorporating Bayesian inference, the decisive factor for how these sources are integrated is their perceived reliability (Friston 2010). More specifically, perceived reliability determines the individual weights assigned to each source of information during the integration process. During FT in our experiment, participants were confronted with at least two sources of sensory cues related to their movements: 1) information that was directly related to their own action, such as forward model outputs, proprioceptive feedback etc.; 2) the experimentally provided explicit visual feedback about their movement direction. The subjective estimate of pointing direction provided by the participants after every trial can be considered a proxy for subjects’ integrated SoA. Given our findings of excessive weighting and adaptation to false visual feedback in patients with OCD, the integration process in these patients was, however, shifted towards “external” sensory information. This occurred, according to Bayesian inference, in a way *as if* “internal” agency cues were less reliable, despite their actual reliability in the domains of motor control and self-action perception in fact was indistinguishable from healthy controls. As previous theories have pointed out, such a deficient integration of internal signals lies at the core of OCD symptomatology (Lazarov, Liberman et al. 2014, Schmidt, Wagner et al. 2021). Accordingly, using a computational model of OCD, based on the Bayesian principles described above, Fradkin and colleagues predicted an *excessive reliance on sensory information* due to patients’ difficulties in predicting changes caused by self-actions (Fradkin, Adams et al. 2020). Similarly, the *seeking proxies for internal states (SPIS) model* suggests that patients with OCD have an *attenuated access* to internal states which makes them prone to rely more strongly on explicit feedback (Lazarov, Liberman et al. 2014). This assumption is supported by studies in which high OC participants or OCD patients exhibited deficits in correctly adjusting muscle tension and rely more strongly on external feedback (Lazarov, Dar et al. 2010, Lazarov, Dar et al. 2012, Lazarov, Liberman et al. 2014) as well as studies demonstrating a diminished interoceptive sensitivity (Schultchen, Zaudig et al. 2019, Demartini, Nisticò et al. 2021). Similarly, Ezrati suggested decreased access to proprioceptive information in high OC individuals to be the reason behind an overreliance on explicit feedback in a reach adaptation paradigm (Ezrati, Sherman et al. 2018). In contrast to this previous work, the patients in our study did not show any such impairment in the perceptual (PT) or motor (MT) domain, which clearly speaks against a general deficit in proprioception or in accessing “internal cues”. Our results rather show that patients can perform and perceive their own actions accurately in situations when no other (possibly conflicting) sources of information are available. We therefore propose that rather than not being able to either access internal states or accurately predict changes brought about by own actions, OCD patients integrate their sensory predictions to a lesser extent, just as if they were unreliable, despite a lack of actual perceptual or motor deficits. This leads to the overreliance on feedback as shown in previous studies. Importantly, the current study adds to these findings by showing the importance of error attribution and how a deficit in this attribution process could contribute to many of the described SoA deficits in OCD.

A remaining question is why patients integrate internal action-related information to a lesser degree. A possible answer can be derived from the predictive coding account of interoception, which assumes that individual perceptions are based on internal working models, so-called priors (Friston, Schwartenbeck et al. 2014). In the case of OCD, the execution of compulsions is associated with intense negative emotions, as well as the urge to resist them, which often fails. Thus, in OCD the priors regarding internal signals may be rooted in beliefs that information from the body is unreliable, resulting in an overreliance on other cues. Interestingly, we found a clear negative correlation between the reliability of all participants’ PPDs and the amount of feedback weighting, which supports the general framework of reliability-based sensory integration. However, the stronger feedback influence in patients cannot be explained by deficient reliability, as we found no group differences in this domain. This is an important distinction to the findings in patients with schizophrenia suffering from delusions of control, who also show increased weighting of external feedback which is most likely based on an unreliable perception of their own actions due to imprecise forward models (Blakemore, Wolpert et al. 2000, Frith, Blakemore et al. 2000, Lindner, Thier et al. 2005, Synofzik, Thier et al. 2010). Future studies are needed to determine whether the enhanced weighting of external cues in OCD is indeed related to particular priors, and, in case, how these priors can be modified via therapy. One recent study has shown interoceptive awareness, i.e., the ability to access bodily signals, was not improved following standard cognitive behavior therapy treatment in OCD patients (Schultchen, Zaudig et al. 2019). Importantly, this treatment approach does not focus on perception of bodily state as a treatment target. Considering the current findings, strengthening the integration of internal agency cues in OCD may be an important treatment target to extend existing intervention approaches.

It is important to note that the current study is also the first to include the experimental manipulation of factors related to outcome valence and social context in the investigation of SoA in self-generated actions in OCD. This was implemented by allowing participants to gain additional money for themselves (self block) or for a charitable foundation (other block). While these experimental manipulations had no influence in PT, there was an influence on FT within and across participant groups. More precisely, there was a significant linear increase across the experimental conditions (baseline < self < other) in the extent to which false visual feedback influenced participants’ perception. Furthermore, when the patient group was divided into two subgroups based on the severity of current OCD symptoms, we found that the more severe the current OCD symptoms of the patients, the more they relied on the strongly rotated visual feedback at the expense of their internal movement information in the other block. In contrast to previous studies however, we found no evidence that – apart from the influence of the social context – the manipulation of valence through monetary gains vs. losses additionally affects the SoA (Takahata, Takahashi et al. 2012, Wilke, Synofzik et al. 2012, Yoshie and Haggard 2013, Tanaka and Kawabata 2019). Previous studies manipulating outcome valence have arrived at conflicting results, with reports suggesting both increased SoA for positive as well as negative outcomes. It is possible that the valence manipulation in the current study was not of sufficient magnitude, especially since gains and losses per trial were on average only .04 Euros. Thus, Increasing the amount of monetary gains or using other modalities, such as pain stimuli, may be a feasible approach to further tackle the influence of valence outcome on SoA in future studies examining agency in OCD. In addition, gains and losses did not depend on participants’ actual performance (compare methods), which also could explain the lack of outcome valence on the SoA.

In our study we found clear evidence for the influence of social factors on SoA, indicating that when self-generated actions had consequences for others, the weighting of external cues was strongest across all participants. This fits to evidence from behavioral economics, suggesting that individuals generally tend to exhibit a more generous giving behavior to counterparts who are perceived as needing help or trigger an empathic response (DeScioli and Krishna 2013, Klimecki, Mayer et al. 2016). Thus, the observed effects are likely due to an enhanced emotional relevance of action outcomes. OCD patients may be more vulnerable to such distortions due to a generally enhanced moral sensitivity and sense of responsibility, thereby increasing the emotional salience when their actions “cause” consequences for social counterparts.

In sum, our study showed increased weighting of false visual feedback regarding self-actions in OCD. Furthermore, this increased weighting was also evident in a trial-by-trial adaptation to this false feedback. Importantly, this deficit was not due to inaccurate or unreliable actions or the perception thereof. Given earlier research demonstrating that the attribution of a perceived movement error to oneself is a prerequisite for adaptation processes, we suggest that this change in the integration of different agency cues speaks in favor of an increased attribution of external errors to oneself in OCD. We propose that this tendency to incorrectly assign sensory information to one’s own making, contributes to many of the SoA distortions described in OCD. The basis for this mechanism could be that patients’ priors give less weight to “internal” information about self-action and thereby lead to an overreliance on other, external action-related feedback. A specific training tailored at increasing perceived reliability and weight of internal action-related information could therefore prove fruitful in ameliorating OCD symptomatology.

## Acknowledgments

This project has been funded by the Fortune-Program (project number 2571-0-0), an intramural funding program of the medical faculty at the University of Tübingen.

We thank Julia Henner for support in data collection.

