## Supplement for "Disturbance of the Sense of Agency in Obsessive-Compulsive Disorder and its Modulation by Social Context"

### Supplemental Information

### Participants

***Table S1 Sample information.*** *OCI-R: Obsessive-Compulsive Inventory-Revised, OCTCDQ: Obsessive-Compulsive Trait Core Dimensions Questionnaire, Y-BOCS: Yale-Brown Obsessive-Compulsive Scale.*

| Patient | Age | Sex | OCI-R | OCTCDQ | Y-BOCS |
|---|---|---|---|---|---|
| **1** | 42 | m | 21 | 57 | 30 |
| **2** | 24 | f | 17 | 30 | 14 |
| **3** | 59 | f | 23 | 63 | 21 |
| **4** | 26 | f | 19 | 27 | 15 |
| **5** | 28 | f | 39 | 64 | 28 |
| **6** | 39 | m | 19 | 39 | 16 |
| **7** | 24 | f | 27 | 60 | 26 |
| **8** | 23 | m | 23 | 44 | 18 |
| **9** | 42 | f | 27 | 47 | 14 |
| **10** | 24 | f | 37 | 50 | 21 |
| **11** | 28 | f | 27 | 44 | 23 |
| **12** | 24 | f | 27 | 45 | 19 |
| **13** | 27 | m | 13 | 37 | 28 |
| **14** | 29 | f | 44 | 61 | 33 |
| **15** | 32 | f | 24 | 10 | 27 |
| **16** | 26 | f | 31 | 54 | 10 |
| **17** | 31 | m | 6 | 16 | 13 |
| **18** | 24 | f | 24 | 41 | 12 |
| **19** | 54 | f | 20 | 20 | 31 |
| **20** | 57 | m | 26 | 21 | 24 |
| **21** | 26 | f | 33 | 66 | 32 |
| **22** | 38 | f | 10 | 27 | 28 |
| **23** | 30 | m | 43 | 59 | 33 |
| **24** | 54 | m | 10 | 21 | 7 |
| **25** | 39 | f | 28 | 56 | 27 |
| **26** | 61 | m | 4 | 5 | 9 |
| **27** | 51 | f | 26 | 19 | 12 |
| **28** | 49 | f | 31 | 30 | 17 |
| **29** | 48 | m | 34 | 35 | 22 |
| **30** | 30 | f | 31 | 50 | 22 |
| **31** | 25 | f | 29 | 56 | 19 |
| **32** | 55 | m | 13 | 8 | 10 |
| **33** | 24 | f | 13 | 16 | 19 |
| **34** | 28 | f | 23 | 47 | 38 |
| **35** | 32 | f | 43 | 70 | 15 |
| **36** | 48 | f | 32 | 30 | 31 |
| **37** | 34 | f | 25 | 42 | 32 |
| **38** | 38 | m | 19 | 61 | 24 |

| Control | Age | Sex | OCI-R | OCTCDQ |
|---|---|---|---|---|
| **1** | 26 | f | 5 | 8 |
| **2** | 28 | f | 6 | 13 |
| **3** | 22 | m | 8 | 8 |
| **4** | 23 | f | 0 | 0 |
| **5** | 23 | f | 7 | 13 |
| **6** | 22 | f | 0 | 2 |
| **7** | 56 | f | 0 | 0 |
| **8** | 23 | f | 2 | 0 |
| **9** | 24 | f | 1 | 0 |
| **10** | 24 | f | 0 | 4 |
| **11** | 25 | m | 4 | 4 |
| **12** | 24 | f | 6 | 4 |
| **13** | 25 | m | 2 | 3 |
| **14** | 38 | f | 5 | 7 |
| **15** | 30 | m | 2 | 2 |
| **16** | 27 | f | 6 | 8 |
| **17** | 28 | f | 7 | 18 |
| **18** | 25 | f | 2 | 1 |
| **19** | 28 | m | 2 | 2 |
| **20** | 28 | f | 6 | 10 |
| **21** | 55 | f | 6 | 5 |
| **22** | 32 | f | 2 | 5 |
| **23** | 26 | f | 5 | 3 |
| **24** | 24 | f | 4 | 10 |
| **25** | 50 | f | 2 | 7 |
| **26** | 25 | m | 6 | 0 |
| **27** | 24 | f | 7 | 4 |
| **28** | 36 | m | 8 | 12 |
| **29** | 35 | f | 6 | 6 |
| **30** | 31 | f | 2 | 5 |
| **31** | 58 | f | 6 | 3 |
| **32** | 30 | m | 5 | 2 |
| **33** | 21 | f | 3 | 1 |
| **34** | 38 | m | 5 | 4 |
| **35** | 37 | f | 7 | 3 |
| **36** | 53 | m | 3 | 6 |
| **37** | 42 | m | 6 | 3 |
| **38** | 52 | m | 4 | 2 |
| **39** | 30 | m | 7 | 15 |

### Offset Correction

In order to minimize interindividual variance we performed an offset correction of FT for every participant by subtracting the value at 0° rotation (i.e. trials with veridical feedback) from all other rotations. This “centers” all participants on zero in case of veridical feedback. To make sure that no differences between groups are lost using this procedure we first checked for differences in 0° trials with a 3 x 2 x 2 mixed-model ANOVA with the factors block (baseline, self, other), outcome (gain, loss), and group (Control, OCD). This revealed no significant main or interaction effects (see Figure S1 for pre-offset-correction data).

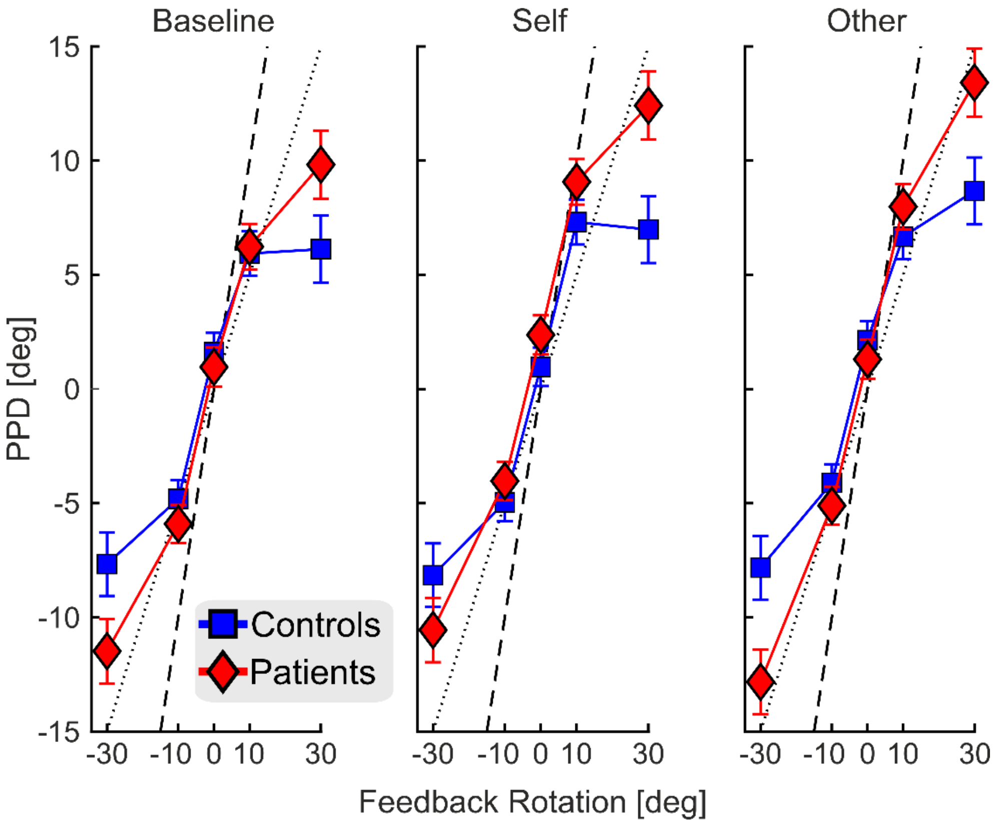


***Figure S1 Results of feedback trials (FT) without offset-correction across experimental blocks and groups (mean ± sem).*** *The ordinate represents the magnitude of PPD (Perceived Pointing Direction) in degrees. Actual rotation of visual feedback is represented on the abscissa. The dotted line (zero PPD) represents the true pointing direction, i.e., participants' perceptual estimates only reflect their actual movement, while the dashed and solid line reflect a perceptual weight of rotated visual feedback, i.e., how strong are participants' perceptual estimates influenced by false visual feedback, by 50% (dashed) or 100% (solid), respectively. Both groups show a bias in the positive, i.e., leftwards, direction in trials with veridical feedback (0°). Repeating the 3 x 2 x 2 mixed-model ANOVA for 30° rotations used in the Results Section with this data (factors block [baseline, self, other], outcome [gain, loss], and group) yields the same general main effects of Group (F(1, 75) = 6.189, p = 0.015) and block (F(2, 150) = 6.964, p =0 .001).*

### Reliability and Feedback Weighting

As detailed in the introduction, the reliability of forward models plays an important role in the construction of a SoA. Therefore, we tested whether the increased weighting of visual feedback in OCD patients could be due to noisier perceptual self-action estimates (as a proxy for forward models). To this end, we compared the reliability, which we defined as the inverse of the variance, of participants' PPD in PT. In order to determine whether the assumed association between perceptual reliability and feedback weighting was present irrespective of subject group, we correlated both variables (i.e. perceptual reliability from PD, PPD at 30° from FT) across all participants using each individual's average reliability across all experimental blocks. We found that PPD reliability was significantly negatively correlated with the strength of feedback weighting at 30° rotation, such that the noisier participants' PPDs were, the stronger the influence of visual feedback on said estimates ($r(75) = -0.424$, $p < 0.001$; Figure S2).

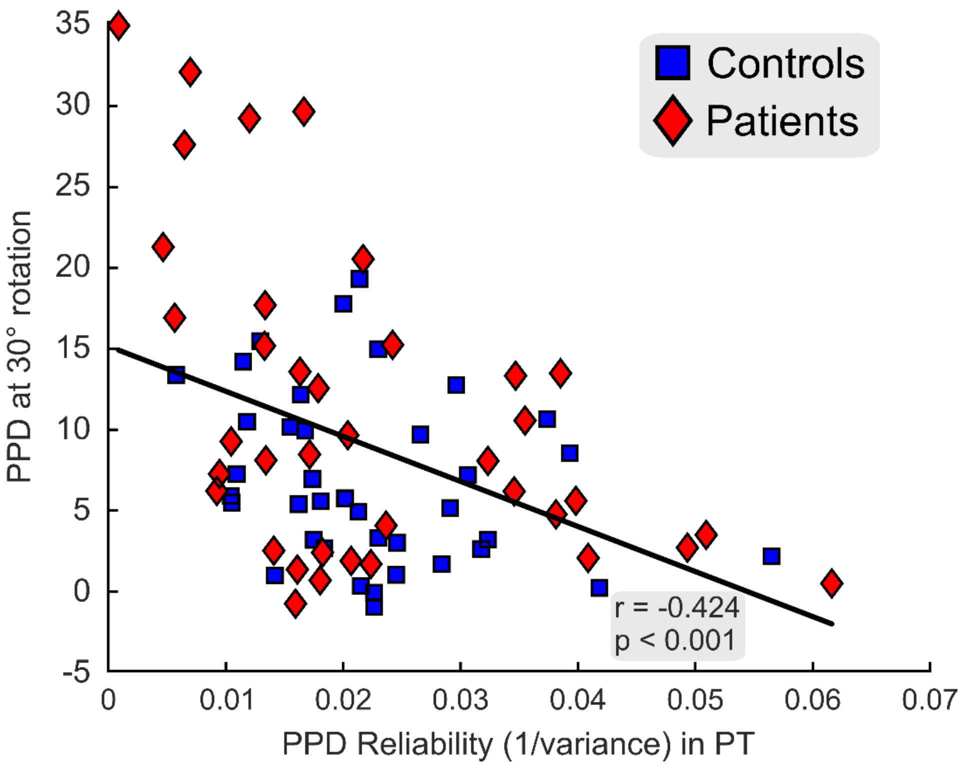


***Figure S2 Correlation between the reliability of participants' perceptual judgments and the perceptual bias in 30° rotated visual feedback.*** *Perceptual reliability was estimated by taking the inverse of PPD variance from PT, i.e., trials in which participants performed pointing movements without visual feedback. This was correlated against average PPDs from FT with*

*30° rotation. PPD reliability was negatively correlated with the strength of feedback weighting, suggesting that the less reliable participants' perceptual judgments were, the stronger they were influenced by false visual feedback.*

### Feedback Weighting and Symptom Strength

In addition to the median split of the patient group according to the score of the OCI-R presented in the main text of this manuscript, we repeated the same analysis with the scores from the OCTCDQ. Figure S3 shows that the OCD group that scored higher in that questionnaire exhibited significantly stronger weighting (Bonferroni corrected) of 30° rotated visual Feedback in the other block relative to baseline (one-sample t-test; $t(18) = 3.307$, $p = 0.004$), while this effect was absent in the OCD group with lower scores (one-sample t-test; $t(18) = 0.096$, $p = 0.925$). A direct comparison between both groups revealed significantly stronger weighting in the high- compared to the low OCD group (independent-samples t-test, $t(36) = -2.184$, $p = 0.036$). Thus, both questionnaires led to the same qualitative results.

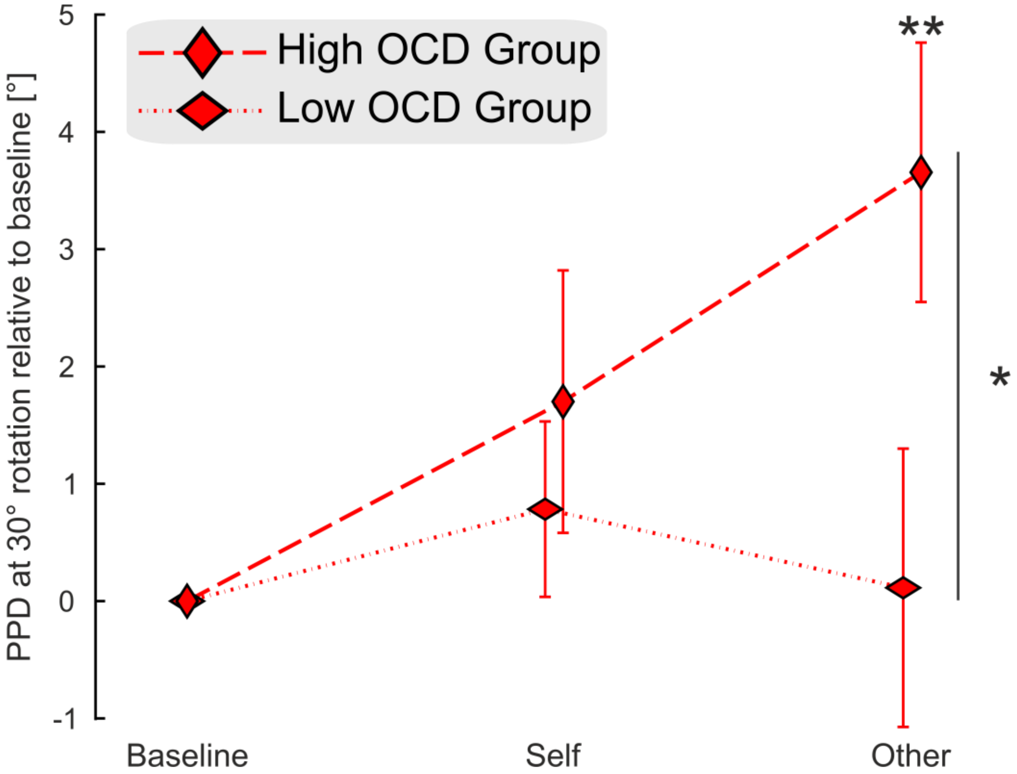


***Figure S3 Differential effect of the experimental block on the two subgroups of patients depending on their symptom severity according to the OCTCDQ.*** *Separating patients into two groups depending on their summed score in the OCTCDQ leads to clear differences in the other block. We expected this block to affect the high OCD group more strongly because it*

*taps into the inflated sense of responsibility and moral sensitivity associated with the OCD symptomatology.* * = $p < 0.05$; ** = $p < 0.005$; *n.s.* = *not significant.*
